# VIDHOP, viral host prediction with Deep Learning

**DOI:** 10.1101/575571

**Authors:** Florian Mock, Adrian Viehweger, Emanuel Barth, Manja Marz

## Abstract

**Motivation:** Zoonosis, the natural transmission of infections from animals to humans, is a far-reaching global problem. The recent outbreaks of Zika virus, Ebola virus and Corona virus are examples of viral zoonosis, which occur more frequently due to globalization. In the case of a virus outbreak, it is helpful to know which host organism was the original carrier of the virus. Once the reservoir or intermediate host is known, it can be isolated to prevent further spreading of the viral infection. Recent approaches aim to predict a viral host based on the viral genome, often in combination with the potential host genome and arbitrarily selected features. These methods have a clear limitation in either the number of different hosts they can predict or the accuracy of their prediction.

**Results:** Here, we present a fast and accurate deep learning approach for viral host prediction, which is based on the viral genome sequence only. To ensure a high prediction accuracy, we developed an effective selection approach for the training data to avoid biases due to a highly unbalanced number of known sequences per virus-host combinations. We tested our deep neural network on three different virus species (influenza A, rabies lyssavirus, rotavirus A). We reached for each virus species an AUG between 0.93 and 0.98, outperforming previous approaches and allowing highly accurate predictions while only using fractions (100-400 bp) of the viral genome sequences. We show that deep neural networks are suitable to predict the host of a virus, even with a limited amount of sequences and highly unbalanced available data. The deep neural networks trained for this approach build the core of the virus-host predicting tool VIDHOP (Virus Deep learning HOst Prediction).

**Availability:** The trained models for the prediction of the host for the viruses influenza A, rabies lyssavirus, rotavirus A are implemented in the tool VIDHOP. This tool is freely available under https://github.com/flomock/vidhop.

**Supplementary information:** Supplementary data are available at DOI 10.17605/OSF.IO/UXT7N

## 1 Introduction

Zoonosis, more specifically, the cross-species transmission of viruses, is a significant threat to human and livestock health because, during a viral breakout, it can be difficult to asses where specific viruses originated (Saez *et al.*, 2015; Longdon *et al.*, 2014). However, this information can be crucial for the effective control and eradication of an outbreak, as the virus needs time to fully adapt to the new human or animal host before it can spread within the new host species. Only with this information, the original host can be separated from humans and livestock. Such isolation can limit the zoonosis and thus can also limit the intensity of a viral outbreak.

Various computational tools for predicting the host of a virus by analyzing its DNA or RNA sequence have been developed. These methods can be divided in three general approaches: supervised learning (Eng *et al.*, 2014; Zhang *et al.*, 2017; Li and Sun, 2018), probabilistic models (Galiez *et al.*, 2017) and similarity rankings (Edwards *et al.*, 2016; Ahlgren *et al.*, 2017). All of these approaches require features with which the input sequence can be classified. The features used for classification are mainly k-mer based on various k sizes between 1-8. In the case of probabilistic models and similarity rankings, not only the viral genomes but also the host genomes have to be analyzed.

Still today, it is mostly unknown how viruses adapt to new hosts and which mechanisms are responsible for enabling zoonosis (Taubenberger and Kash, 2010; Villordo *et al.*, 2015; Longdon *et al.*, 2014). Because of this incomplete knowledge, it is likely to choose inappropriate features, *i.e.*, features of little or no biological relevance, which is problematic for the accuracy of machine learning approaches. In contrast to classic machine learning approaches, deep neural networks can learn features necessary for solving a specific task by themselves.

In this study, we present a novel approach, using deep neural networks, to predict viral hosts by analyzing either the whole or only fractions of a given viral genome. We selected three different virus species as individual datasets for training and validation of our deep neural networks. These three datasets consist of genomic sequences from influenza A, rabies lyssavirus and rotavirus A, composed of 49, 19, and 6 different known host species, respectively. These known viral hosts are often phylogenetically close related. Previous prediction approaches have combined single species or even genera to higher taxonomical groups to reduce the classification complexity to the price of prediction precision (Zhang *et al.*, 2017; Galiez *et al.*, 2017). In contrast, our approach is capable of predicting at the host species level, providing much higher accuracy and usefulness of our predictions.

Our training data consists of at least 100 genomic sequences per virus-host combination. The amount of sequences per combination is unbalanced, which means that some classes are much more common than others. We provide an approach to handle this problem by generating a new balanced training set at each training circle.

Typically the training of recurrent neural networks on very long sequences is very time consuming and inefficient. Truncated backpropagation trough time (TBPTT) (Puskorius and Feldkamp, 1994) tries to solve this problem by splitting the sequences into shorter fragments. We provide a method to regain prediction accuracy lost through this splitting process, leading to fast, efficient learning of long sequences on recurrent neural networks.

In conclusion, our deep neural network approach is capable of predicting far more complex classification problems than previous approaches (Eng *et al.*, 2014; Kapoor *et al.*, 2010; Zhang *et al.*, 2017; Galiez *et al.*, 2017; Li and Sun, 2018). Meaning, it is more accurate for the same amount of possible hosts and can predict for more hosts with similar accuracy. Furthermore, our approach does not require any host sequences, which can be helpful due to the limited amount of reference genomes of various species, even ones that are typically known for zoonosis such as ticks and bats (Dilcher *et al.*, 2012; **?**; Teeling *et al.*, 2018; Van Zee *et al.*, 2007).

## 2 Methods

### 2.1 General workflow

We designed our general workflow to achieve multiple goals:

(**I**) select, preprocess and condense viral sequences with as little information loss as possible (**II**) correctly handle highly unbalanced datasets to avoid bias during the training phase of the deep neural networks (**III**) present the output in a clear, user-friendly way while providing as much information as possible.

Our workflow for creating the deep neural networks, used in VIDHOP, to predict viral hosts consisted of five major steps (see Figure 1). First, we collected all nucleotide sequences of influenza A, rabies lyssavirus, and rotavirus A with a host label from the European Nucleotide Archive (ENA) database (Leinonen *et al.*, 2010). We curated the host labels using the taxonomic information provided by the National Center for Biotechnology Information (NCBI), leading to standardized scientific names for the virus taxa and host taxa. Standardization of taxa names enables swift and easy filtering for viruses or hosts on all taxonomic levels. Next, we divide the sequences from the selected hosts and viruses into three sets: the training set, the validation set, and the test set. We provide a solution to use all sequences of an unbalanced dataset without biasing the training in terms of sequences per class while limiting the memory needed to perform this task. Then, the length of the input sequences is equalized to the 0.95 quantile length of the sequences and subsequently further truncated in shorter fragments and parsed into numerical data to facilitate a swift training phase of the deep neural network. After the input preparation, the deep neural network predicts the hosts for the subsequences of the originally provided viral sequences. In the final step, the predictions of the subsequences are analyzed and combined to a general prediction for their respective original sequences.

**Figure 1:**
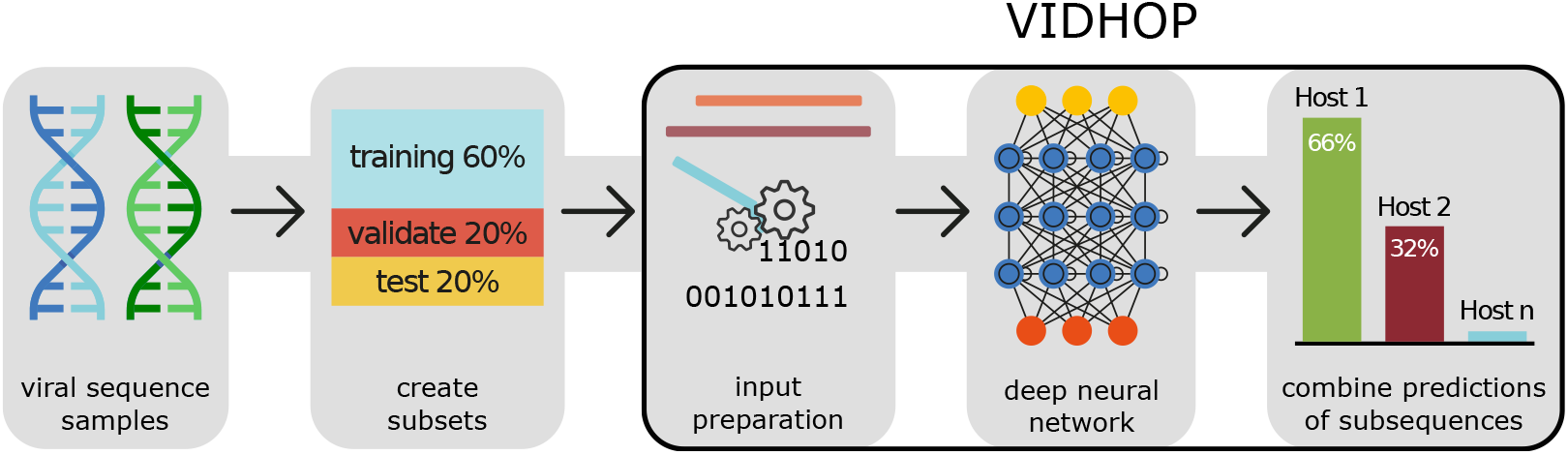
The general workflow consists of several steps. First, suitable viral sequences have to be collected and standardized. Next, these sequences will be distributed into the training set, validation set, and test set. The sequences are then adjusted in length and are parsed into numerical data, which is then used to train the deep neural network. The neural network predicts the host for multiple subsequences of the original input. The subsequence predictions are then combined to a final prediction. The workflow used in VIDHOP to predict new sequences is framed in black.

### 2.2 Collecting sequences and compiling datasets

Accession numbers of influenza A, rabies lyssavirus, and rotavirus A were collected from the ViPR database (Northrop Grumman Health IT and Technologies, 2017) and Influenza Research Database (for Biotechnology Information, 2017) and all nucleotide sequence information were then downloaded from ENA (status 2018-07-12). From the collected data, we created one dataset per virus species, with all known hosts that had at least 100 sequences. All available sequences were used for each of these hosts (see Figure 2).

**Figure 2:**
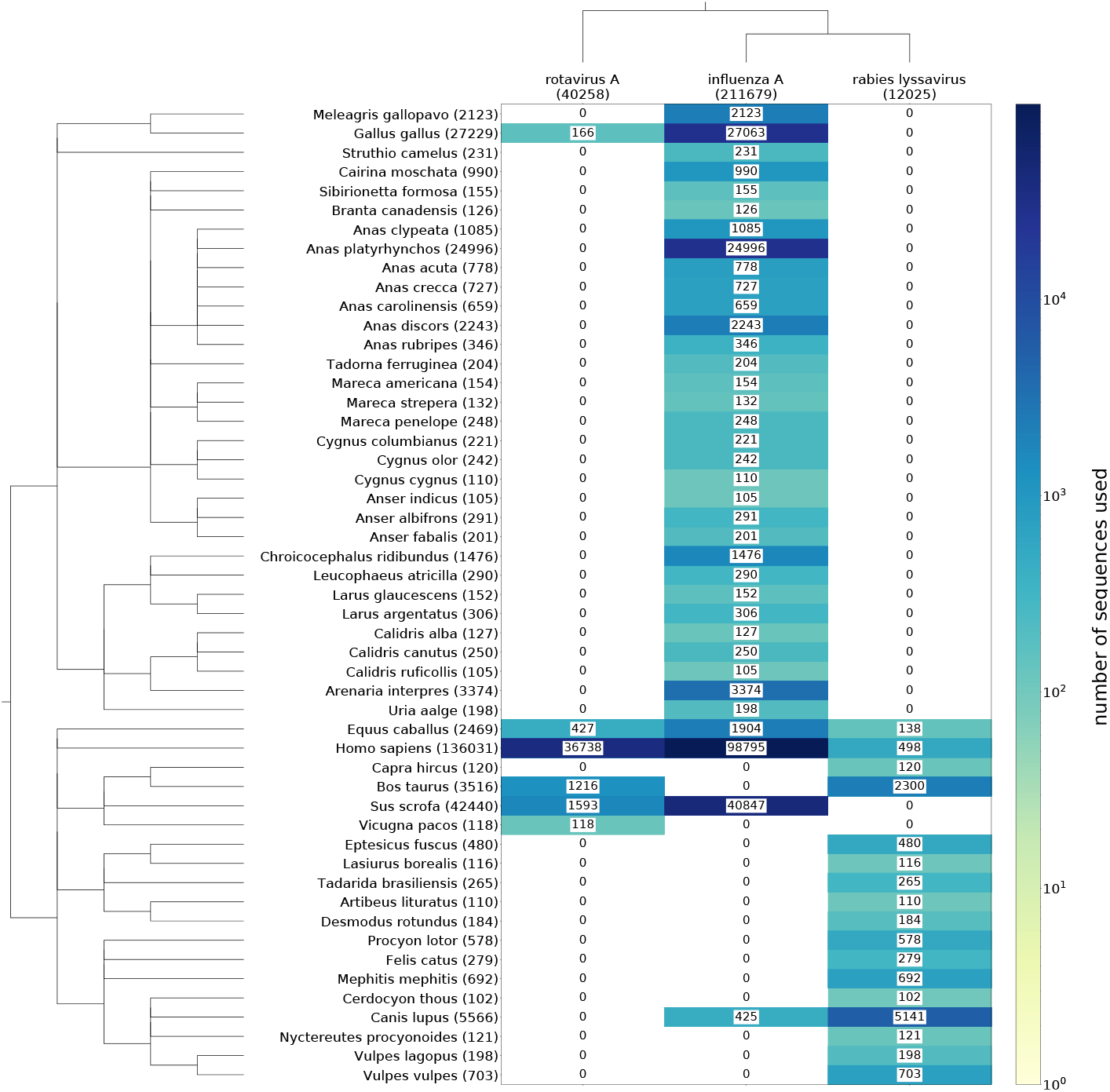
At the top, the examined virus species are listed with the total amount of used nucleotide sequences shown in brackets. On the left, the potential host species are listed together with the total number of available sequences for these three viruses. The matrix lists the corresponding number of sequences for each virus-host combination used in this study. Dendrograms indicate the phylogenetic relationships of the investigated viruses and host species. As it can be seen, there is a clear imbalance in available viral sequences per virus-host pair.

To train the deep neural network, we divided each dataset into three smaller subsets, a training set, a validation set, and a test set. Classically, in neuronal network approaches, the data is divided into a ratio of 60% training set, 20% validation set, and 20% test set with a balanced number of data points per class.

Since in our example, nucleotide sequences are the data points, and the different hosts are the classes, this would lead to an unbiased training per host. But for heavily unbalanced datasets, such as typical viral datasets, the majority of sequences would not be used. This is because the host with the smallest number of sequences would determine the maximum usable amount of sequences per host (see Figure 3).

**Figure 3:**
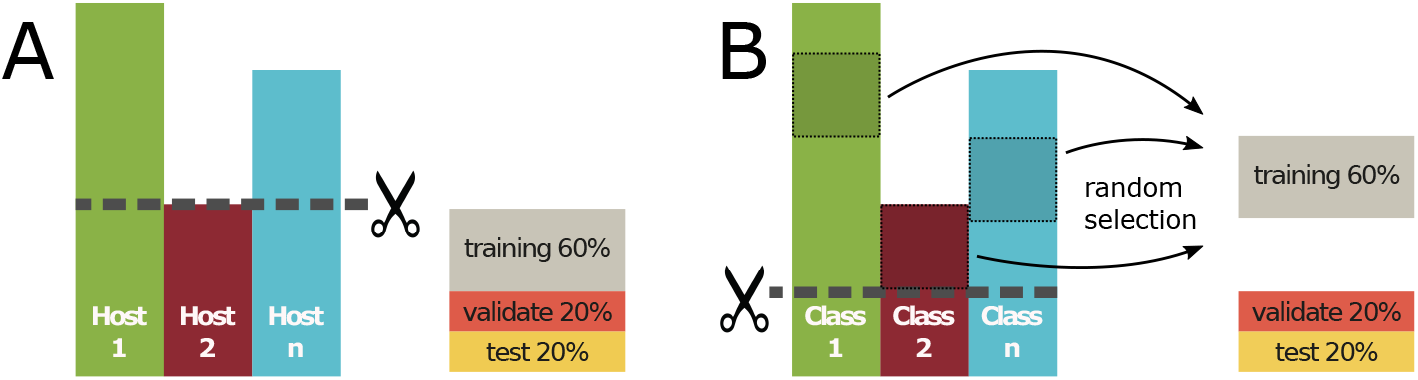
Comparison between the classic approach (**A**) of creating a balanced dataset and the repeated random undersampling (**B**). In the classic approach, the class with the smallest number of data points defines the number of usable data points for all classes. The repeated random undersampling creates every epoch a new random composition of training data points from all data points which are not included in any of both fixed validation set and test set. For every epoch, the training set is balanced according to the data points per class. The repeated random undersampling can use all available data in an unbalanced dataset, without biasing the training set in terms of data points per class, while limiting the amount of computer memory needed.

A more appropriate approach to deal with large unbalanced datasets is to define a fixed validation set and a fixed test set and create variable training sets from the remaining unassigned sequences. In the following, we call this the *repeated random undersampling*. For each training circle (epoch), a new training set is created by randomly selecting the same number of unassigned sequences per host. The number of selected sequences per host corresponds to the number of unassigned sequences of the smallest class. Repeated random undersampling avoids bias in the training set in terms of sequences per host while using all available sequences. Especially hosts with large quantities of sequences benefit from the generation of many different training sets with random sequence composition.

### 2.3 Input preparation

The training data needs to fulfill several properties to be utilizable for neural networks. The input (here, nucleotide sequences) has to be of equal length and also has to be numerical. To achieve this, the length of the sequences was limited to the 0.95 quantile of all sequence lengths by truncating the first positions or in the case of shorter sequences by extension. For sequence extension different strategies were tested and evaluated (see Figure 4, Supplement Table S2):

**Figure 4:**
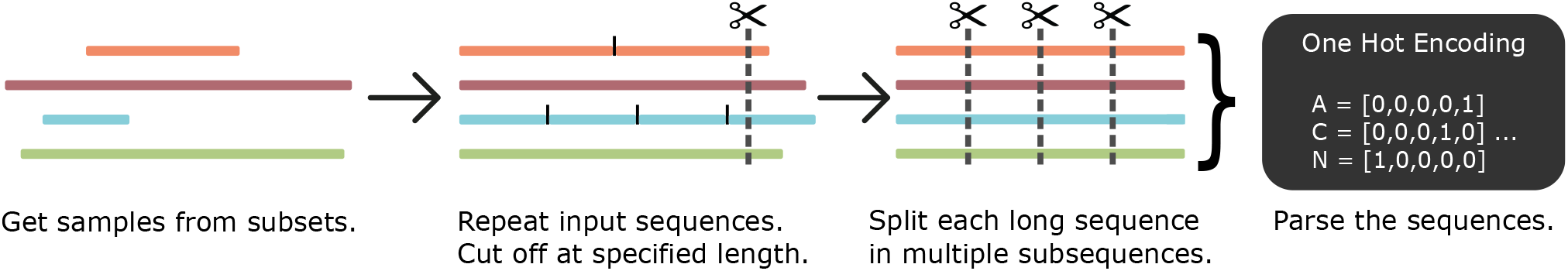
Conversion of input sequences into numerical data of equal length using the *normal repeat* method. Each sequence is extended through self-repetition and is then trimmed to the 0.95 quantile sequence length. Sequences are then split into multiple non-overlapping subsequences of equal length. Each subsequence is then converted via *one hot encoding* into a list of numerical vectors.

- *Normal repeat:* repeats the original sequence until the 0.95 quantile of all sequence lengths is reached, all redundant bases are truncated.
- *Normal repeat with gaps:* between one and ten gap symbols are added randomly between each sequence repetition.
- *Random repeat:* appends the original sequence with slices of the original with the same length as the original. For this purpose, the sequence is treated as a circular list. In a circular list, the end of the sequence is followed by the beginning of the sequence.
- *Random repeat with gaps:* like *random repeat*, but repetitions are separated randomly by one to ten gap symbols.
- *Append gaps:* adds gap symbols at the end of the sequence until the necessary length is reached.
- *Online:* uses the idea of *Online Deep Learning* (Sahoo *et al.*, 2017), i.e., slight modifications to the original training data are introduced and learned by the neuronal network, next to the original training dataset. In our case, randomly selected subsequences of the original sequences are provided as training input. Therefore more diverse data is provided to the neural network.
- *Smallest:* all sequences are cut to the length of the shortest sequence of the dataset.

After applying one of the mentioned input preparation approaches, each sequence is divided into multiple non-overlapping subsequences (see Figure 4). The length of these subsequences ranged between 100 and 400 nucleotides, depending on which length results in the least redundant bases. Using subsequences of a distinct shorter length than the original sequences is a common approach in machine learning to avoid inefficient learning while training long short-term memory networks on very long sequences, see *Truncated Backpropagation Through Time* approach (Sutskever, 2013).

Finally, all subsequences are encoded numerically, using *one hot encoding* to convert the characters A, C, G, T, -, N into a binary representation (*e.g., A* = [1, 0, 0, 0, 0], *T* = [0, 0, 0, 1, 0],− = [0, 0, 0, 0, 0]). Other characters that may occur in the sequence data were treated as the character N.

### 2.4 Deep neural network architecture

Its underlying architecture dramatically determines the performance of the neural network. This architecture needs to be complex enough to use the available information fully but, at the same time, small enough to avoid overfitting effects.

All our models (i.e., the combination of the network architecture and various parameter such as the optimizer or validation metrics) were built with the `Python` (version 3.6.1) package `Keras` (Chollet *et al.*, 2015) (version 2.2.4) using the `Tensorflow` (Abadi *et al.*, 2015) (version 1.7) back-end.

In this study, two different models were built and evaluated to predict viral hosts only given the nucleotide sequence data of the virus (see Figure 5). The architecture of our first model consists of a three bidirectional LSTM layers (Hochreiter and Schmidhuber, 1997), in the following referred to as *LSTM* architecture (see Figure 5 A). This bidirectional LSTM tries to find longterm context in the input sequence data, presented to the model in forward and reverse direction, which helps to identify interesting patterns for data classification. The LSTM layers are followed by two dense layers were the first collects and combines all calculations of the LSTMs, and the second generates the output layer. Each layer consists of 150 nodes with an exception to the output layer, which has a variable number of nodes. Each node of the output layer represents a possible host species. Since each tested virus dataset contains different numbers of known virus-host pairs, the number of output nodes varies between the different virus datasets. This architecture is similar to those used in text analysis but specifically adjusted to handle long sequences, which are typically problematic for deep learning approaches.

**Figure 5:**
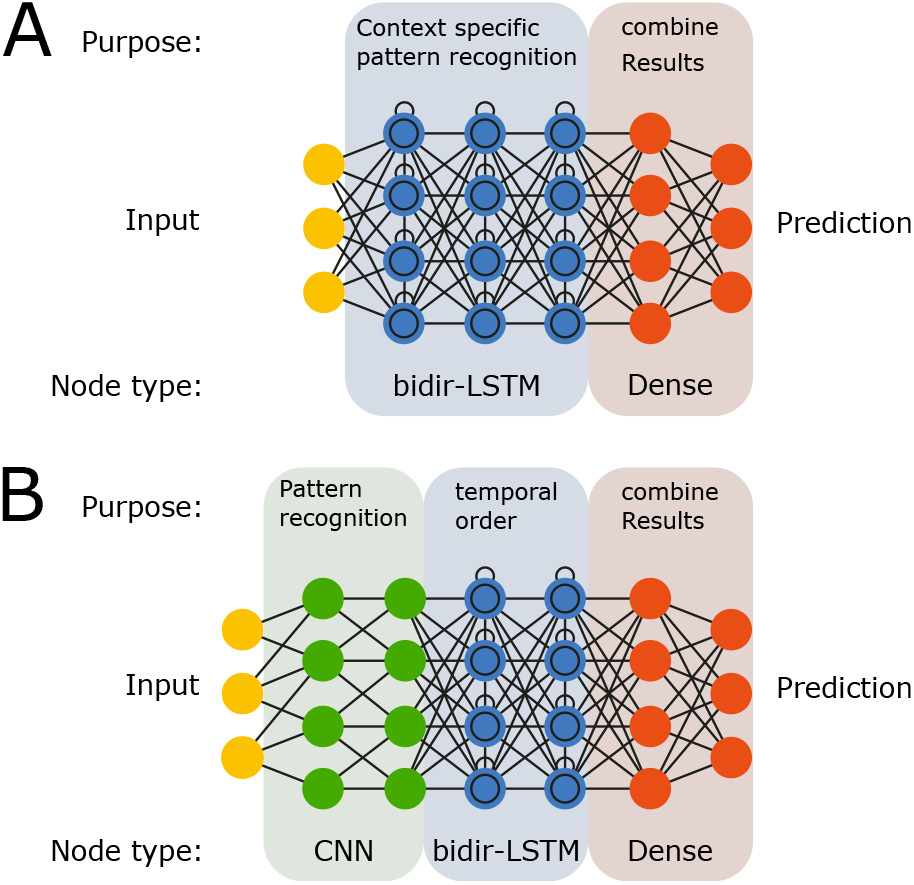
Comparison of the two evaluated architectures. The first architecture (**A**) is similar to neural networks for text analysis. The bidirectional LSTM analyzes the sequence forwards and backward for meaningful patterns, having an awareness of the context as it can remember previously seen data. This architecture is a classic approach for analyzing sequences with temporal information, like literature text, stocks, weather. The second architecture (**B**) uses CNN nodes, which are common in image recognition, to identify meaningful patterns and combines them into complex features that can then be used by the bidirectional LSTM layers. This architecture is typically used in either more unordered data, such as images, or data with more noise, such as the base-caller output of nanopore sequencing devices (Teng *et al.*, 2018).

The second evaluated architecture uses two layers of *convolutional neural networks* (CNN) nodes, followed by two bidirectional LSTM layers and two dense layers. In the following we will refer to this as the *CNN+LSTM architecture* (see Figure 5 B). Similar to the LSTM architecture, each layer consists of 150 nodes with an exception to the output layer. The idea behind this architecture is that the CNN identifies important sequence parts (first layer), combines the short sequence features to more intricate patterns (second layer), which can then be put into context by the LSTMs, which can remember previously seen patterns.

### 2.5 Deep neural network training

The training was done using the *repeated random undersampling* as described above (see Figure 3 B), *i.e.*, having a fixed validation set and test set while the training set was newly compiled during each epoch. All classes had an equal amount of sequences during each epoch. This balancing step results in an equal likelihood to observe each class while training, eliminating the bias of unbalanced training sets. Both neural networks were trained for 500 epochs during all performed tests. After each epoch, the quality of the model was evaluated by predicting the hosts of the validation set, comparing the prediction with the true known virus-host pairs. As metrics, the accuracy and the categorical cross-entropy were used. If the current version of the model performed better, *i.e.*, it had a lower validation loss, or higher validation accuracy than in previous epochs, the weights of the network were saved.

After training, the model weights with the lowest validation loss and model weights with the highest validation accuracy were applied for predicting the test set.

### 2.6 Final host prediction from subsequence predictions

When given a viral nucleotide sequence, the neural network returns the activation score of the corresponding output nodes of each host. The activation scores of all output nodes add up to 1.0 and can, therefore, be treated as probabilities. Thus, the activation score of each output node represents the likelihood of the corresponding species to serve as a host of the given virus sequence. Due to the splitting of the long sequence into multiple subsequences (see Figure 4), the neural network predicts potential hosts for every subsequence. The predictions of the subsequences are then combined to the final prediction of the original sequence. Several approaches to combine the subsequence predictions into a final sequence prediction were evaluated (for a detailed example see supplement S1):

1. *Standard:* shows the original accuracy for each subsequence.
2. *Vote:* uses a majority vote on all subsequences to determine the prediction.
3. *Mean:* calculates the mean activation score per class on all subsequences and predicts the class with the highest mean activation.
4. *Standard deviation:* similar to *Mean* but weights each subsequence with its standard deviation. Subsequences with more distinct predictions get a higher weight.

After combining the subsequence predictions, the single most likely host can be provided as output. However, this limits the prediction power of the neural network. For example, a virus that can survive in two different host species will likely have a high activation score for both hosts. Our tool VIDHOP reports all possible hosts that reach a certain user-defined likelihood, or it can report the *n* most likely hosts, where *n* is also a user-adjustable parameter.

## 3 Discussion

To evaluate our deep learning approach, we applied it to three different datasets, each containing a great number of either influenza A, rabies lyssavirus or rotavirus A genome sequences, and the respectively known host species. We tested two different architectures and seven different input sequence preparation methods. For all fourteen combinations, a distinct model was trained for a maximum of 1500 epochs. The training was stopped early if the validation accuracy did not increase in the last 300 epochs. For each combination, the prediction accuracy was tested using none or any of the described subsequence prediction approaches.

### 3.1 Rotavirus A dataset

The rotavirus A dataset consists of over 40,000 viral sequences, which are associated with one of six phylogenetically distinct host species. Six different hosts result in an expected random accuracy of *∼*16.67 %. Both tested architectures achieve very high prediction accuracies, even for 239 nucleotide long subsequence (see Table 1). The prediction accuracy is influenced not only by the architecture but especially by the input preparation strategy. Overall, the CNN+LSTM architecture achieves higher accuracy than the LSTM architecture with 85.28 %, and 82.88 %, respectively. The highest accuracy was observed with the combination of the CNN+LSTM architecture and the *online* input preparation. Note that the LSTM architecture has difficulties in learning with some input preparation methods (see LSTM and *online*). On the one hand, this is probably due to the relatively long input sequence since the LSTM must propagate the error backwards through the entire input sequence and update the weights with the accumulated gradients. The accumulation of gradients over hundreds of nucleotides in the input sequence may cause the values to shrink to zero or result in inflating values (Werbos *et al.*, 1990; Tallec and Ollivier, 2017). On the other hand, the variability of the online input could enhance the difficulty of finding working features.

**Table 1.** Host prediction accuracy in percent on the rotavirus A, rabies lyssavirus, and influenza A dataset with different architectures and input preparation strategies. The input preparation strategy with the highest accuracy for each architecture is marked dark grey, the second-best in light grey. Expected accuracy by chance is *∼*16.67 % for rotavirus A, *∼*5.88 % for rabies lyssavirus and *∼*2.78 % for influenza A.

| Input preparation<br>vs training setup | normal<br>repeat | normal<br>repeat gaps | random<br>repeat | random<br>repeat gaps | append<br>gaps | online | smallest |
| --- | --- | --- | --- | --- | --- | --- | --- |
| LSTM *rotavirus A* | 78.64 | 82.88 | 78.86 | 82.12 | 43.18 | 25.14 | 75.83 |
| CNN+LSTM *rotavirus A* | 80.38 | 83.79 | 80.30 | 83.03 | 43.78 | 85.28 | 72.50 |
| LSTM *rabies lyssavirus* | 73.68 | 72.37 | 72.98 | 74.26 | 24.39 | 9.39 | 69.41 |
| CNN+LSTM *rabies lyssavirus* | 74.24 | 73.82 | 74.39 | 72.82 | 24.53 | 72.81 | 73.82 |
| LSTM *influenza A* | 48.08 | 50.14 | 48.52 | 49.40 | 35.21 | 49.75 | 48.08 |
| CNN+LSTM *influenza A* | 48.38 | 49.28 | 47.22 | 49.40 | 43.39 | 48.36 | 46.94 |

With prediction accuracies over 82%, on the subsequence level, both the LSTM and the CNN+LSTM architectures indicate that they can identify meaningful classification features. The main differences between the two architectures in the prediction accuracies derive from the applied input preparation strategy. In total, the host prediction quality of rotavirus A sequences achieves an area under the curve (AUC) of 0.98 (see Supplement Figure S3). A high AUC is not unsuspected since it is known that rotavirus A shows a distinct adaptation to their respective host (Martella *et al.*, 2010).

### 3.2 Rabies lyssavirus dataset

The rabies lyssavirus dataset consists of more than 12,000 viral sequences, which are associated with 17 different host species, including closely and more distantly related species. This results in an expected random accuracy of *∼*5.88 %. Despite using only a subsequence length of 100 bases, the accuracy of each subsequence prediction is very high (see Table 1). The LSTM and the CNN+LSTM architecture reach very similar accuracies with 74.26 %, and 74.39 %, respectively. The differences in prediction accuracy between the architectures per input preparation method are small. An exception from this is the *online* input preparation method. Similar to the rotavirus A dataset, the LSTM is not able to train well when using the *online* input preparation method. The highest accuracy per subsequence is reached with the combination of the *random repeat* input preparation and the CNN+LSTM architecture. Compared to the rotavirus A dataset, the higher amount of host species and closer relation between them makes the rabies lyssavirus dataset harder to predict. In total, the host prediction quality of rabies lyssavirus sequences achieves an AUC of 0.98 (see Supplement Figure S4).

### 3.3 Influenza A dataset

The influenza A dataset is with more than 213,000 viral sequences and 36 associated possible host species (32 of them are closely related avian species), the most complex of the three evaluated datasets. With 36 different hosts, the expected random accuracy is *∼*2.78 %, which we greatly exceed with an accuracy of over 50 %. The predictions based on 400 nucleotide long influenza A subsequences reached comparable accuracies for nearly all input preparation methods, one notable exception was *append gaps*, (see Table 1). Unlike for the rotavirus A and rabies lyssavirus dataset, the LSTM architecture outperforms the CNN+LSTM architecture with 50.14 %, and 49.40 % host prediction accuracy. Nonetheless, the differences in prediction accuracy between the architectures per input preparation method are again small. The overall best-performing variant is a combination of the LSTM architecture with the *normal repeat gaps* input preparation.

The deep neural network achieved an AUC of 0.94 (see supplement Figure S5). Despite the close evolutionary distance between the given host species, the trained neuronal network was able to identify potential hosts accurately. We assume that some of the influenza A viruses which are part of the investigated dataset are capable of infecting not only one but several host species, *i.e.*, a single viral sequence can occur in more than one host. However, since we only consider a single host species for each tested viral sequence within the test set, the measured accuracy is most likely an underestimation.

### 3.4 Best practice and useful observations

Overall, the host prediction quality for short subsequences for all three datasets is very high, indicating that an accurate prediction of a viral host is possible even if the given viral sequence is only a fraction of the corresponding genome’s size. Both architectures are suitable for host prediction, but the more complex the prediction task and the data set, the more favorable the LSTM appears. Nevertheless, for fast prototyping, it makes sense to use CNN+LSTM as it trains around four times faster and reaches comparable results. Furthermore, the CNN+LSTM architecture showed no difficulty in learning long input sequences (see Supplement Figure S1). In contrast, the LSTM architecture frequently remained in a state of random accuracy for a long time during training (see Supplement Figure S2).

We observed *random repeat gaps* and *normal repeat gaps* to be the most suited input preparations for the LSTM architecture, as they achieved the highest accuracies. The final selection of the best working input preparation seems to depend on the virus species.

The *random repeat gaps* approach provides the neural network with an almost random selection of the original sequence. All selections are separated by gaps. The first subsequences are always the beginning of the original sequence, whereas the last subsequences consist of random selections.

With the *normal repeat gaps* approach, the neural network can identify not just the start but also the end of the original sequence because the ends are marked by gaps. This may provide a useful context detection for the LSTM layer.

A completely random selection, as in the *online* approach, seems to be too diverse. Non-random approaches such as *normal repeat* seem to lead to faster overfitting of the training set, thus limiting the ability of the deep neural network to identify general usable features.

*Smallest* and *append gaps* proved to be unsuitable methods for input preparation. Here *append gaps* leads to a prediction of subsequences without usable information because they consist only of gaps, whereas *smallest* limits the available information so much that a prediction becomes inaccurate.

### 3.5 Combining subsequence host predictions results in higher accuracy

Among the tested combination approaches of the subsequence-predictions, *std-div* was observed to perform best (see Table 2). With the combination of all subsequence predictions, the accuracy rises between 2.2 %– 4.6 %, with a mean increase of 3.2 %. This result shows that a combination of the host prediction results of all subsequences of a given viral sequence can increase the overall prediction accuracy. Presumably, the prediction combination approaches can compensate for the possible information loss caused by the sequence splitting process during the input preparation.

**Table 2.** Percentage accuracy of the best working input preparation with respect to the combination methods of the subsequence predictions. The combination method with the highest accuracy for each combination is marked in grey.

| Combination method<br>vs training setup | Standard | Voting | Mean | Std-div |
| --- | --- | --- | --- | --- |
| LSTM *rotavirus A*, normal repeat gaps | 82.88 | 87.50 | 86.67 | 87.50 |
| CNN+LSTM *rotavirus A*, online | 85.28 | 87.50 | 87.50 | 88.33 |
| LSTM *rabies lyssavirus*, random repeat gaps | 74.26 | 76.47 | 76.18 | 76.18 |
| CNN+LSTM *rabies lyssavirus*, random repeat | 74.39 | 77.06 | 76.18 | 76.47 |
| LSTM *influenza A*, normal repeat gaps | 50.14 | 52.92 | 54.31 | 54.31 |
| CNN+LSTM *influenza A*, random repeat gaps | 49.40 | 52.64 | 52.22 | 52.64 |
| Mean accuracy | 69.40 | 72.35 | 72.18 | 72.57 |

### 3.6 VIDHOP outperforms other approaches

Besides evaluating our deep learning approach on the three virus datasets, we compared our results with a similar study. Our approach predicts hosts on the species level, whereas most other studies are limited to predicting the host genera (Zhang *et al.*, 2017; Galiez *et al.*, 2017) or even higher taxonomic groups (Eng *et al.*, 2014; Kapoor *et al.*, 2010).

In a relatively comparable study, Le *et al.* (Li and Sun, 2018) also tried to predict potential hosts on species level for influenza A and rabies lyssaviruses. In their study, they mainly tested three different approaches, which were mostly combinations of already published methods (Ahlgren *et al.*, 2017; Zhang *et al.*, 2017; Kapoor *et al.*, 2010), including a support vector machine approach and two sequence similarity approaches, one of which alignment-based, the other without alignment. These three methods represent the state of the art.

The rabies lyssavirus dataset from Le *et al.* consisted of 148 viruses and 19 associated bat host species. Our rabies lyssavirus dataset consists of 12,025 viruses and has 17 associated host species, but none of them is a bat species. Since both data sets do not have a common species, the difficulty of both prediction tasks is difficult to estimate when compared to each other, and therefore comparability is limited. Le *et al.* reached an accuracy below 79 % on the bat species dataset with the similarity-based approaches, one of which relies on alignment data and *vice versa*. The SVM reached an accuracy below 76 %. On our dataset, VIDHOP reached a similar accuracy of around 77 %. However, in contrast to our analysis, Le *et al.* used an unbalanced dataset, which often leads to an overestimation of the prediction accuracy. Furthermore, the presented accuracy from Le *et al.* is based on n-fold cross-validation. N-fold cross-validation lowers the comparability with other studies since the quasi-standard for accuracy determination is 10-fold cross-validation. When applying 10-fold cross-validation, their host prediction accuracy for their rabies lyssavirus dataset drops under 65 %.

The influenza A dataset from Le *et al.* consisted of 1,200 viral sequences and six associated host species. For this dataset, they reached a host prediction accuracy of below 61 % with the alignment-free method (below 40 % when applying 10-fold cross-validation). The alignment-based method reached bellow 52 %, and the SVM bellow 57 %. Our influenza A dataset consists of 211,679 viral sequences and has 36 associated host species, including the six species from the Le *et al.* dataset. Our deep learning approach reached a host prediction accuracy of 54.31 %, which is very good, given that we had to predict six times the number of potential host species with closer phylogenetic relationships among them. To reach better comparability by considering the number of classes, we calculated the average accuracy (see Equation 1).

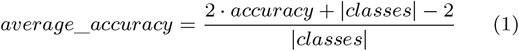

With an average accuracy of 97.46 % VIDHOP surpassed the previous methods which reached an average accuracy between 84 % and 87 % (see Table 3).

**Table 3.** Comparison of VIDHOP with previous approaches. Due to differences in the number of predicted hosts (VIDHOP 36 hosts, Le *et al.* 6 hosts), the average accuracy (Sokolova and Lapalme, 2009) was chosen for comparison. The prediction method with the highest accuracy is marked in grey.

| Average accuracy comparison | *influenza A* |
| --- | --- |
| VIDHOP | 97.46 |
| Le et al. alignment free method | 87.00 |
| Le et al. alignment based method | 84.00 |
| Le et al. SVM | 85.67 |

To our best knowledge, no comparable study exists for the rotavirus A dataset.

VIDHOP reached on all three datasets a very high average accuracy between 96.11 % for rotavirus A, 97.30 % for rabies lyssavirus and 97.46 % for influenza A (see Table 4). These results indicate the versatility of the presented deep learning approach for the task of host prediction.

**Table 4.**
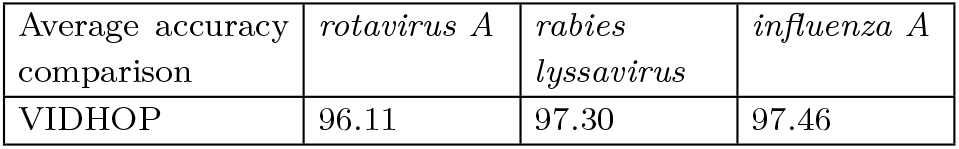
Comparison of the average accuracy of VIDHOP on the three different datasets.

## 4 Conclusion

In this study, we presented the tool VIDHOP and investigated the usability of deep learning for the prediction of hosts for distinct viruses, based on the viral nucleotide sequences alone. We established a simple but very capable prediction pipeline, including possible data preparation steps, data training strategies, and a suitable deep neural network architecture. Besides, we provide three different neural network models, which can predict potential hosts for either influenza A, rotavirus A, or rabies lyssavirus, respectively. These deep neural networks are used in VIDHOP and use genomic fragments shorter than 400 nucleotides to predict potential virus hosts directly on a species level. In contrast to similar approaches, this is a more complex task than performing host prediction only on the genera level (Zhang *et al.*, 2017; Galiez *et al.*, 2017) or even higher taxonomic groups (Eng *et al.*, 2014; Kapoor *et al.*, 2010). Moreover, our approach can predict more hosts with comparable accuracy than previous approaches. The consistently high average accuracy of VIDHOP, on all three datasets, indicates the versatility of the deep learning approach we used.

Additionally, we addressed multiple problems that arise when using DNA or RNA sequences as input for deep learning, such as unbalanced datasets for training and the problem of inefficient learning of recurrent neural networks (RNN) on long sequences. We evaluated different solutions to solve these problems and observed that splitting of the original virus genome sequence in combination with merging the prediction results of the generated subsequences leads to fast and efficient learning on long sequences. Furthermore, the use of unbalanced datasets is possible if a new balanced training set is generated by repeated random undersampling (a random selection of available sequences) for every single epoch during the training phase.

With the use of deep neural networks for host predicting of viruses, it is possible to rapidly identify the host, without the use of arbitrarily selected learning features, for a large number of host species. This allows us to identify the original host of zoonotic events and makes it possible to swiftly limit the intensity of a viral outbreak by separating the original host from humans or livestock.

In future approaches, it could be interesting to investigate the use of newly developed deep neural network layers, such as transformer self-attention layers (Vaswani *et al.*, 2017). This layer type has been shown to perform well with character sequences (Al-Rfou *et al.*, 2018), such as DNA or RNA sequences, potentially allowing for a further increase in the prediction quality.

## Author Contributions

*Conceptualization: FM, AV and MM*. *Data curation: FM*. *Formal analysis: FM*. *Data interpretation: FM*. *Methodology: FM, AV and MM*. *Validation: FM*. *Visualization: FM*. *Writing – Original Draft Preparation: FM*. *Writing – Review & Editing: EB*. *Project Administration: MM*. *Supervision: MM*. *Funding acquisition: MM*.

## Supplementary Information

**Figure S1:**
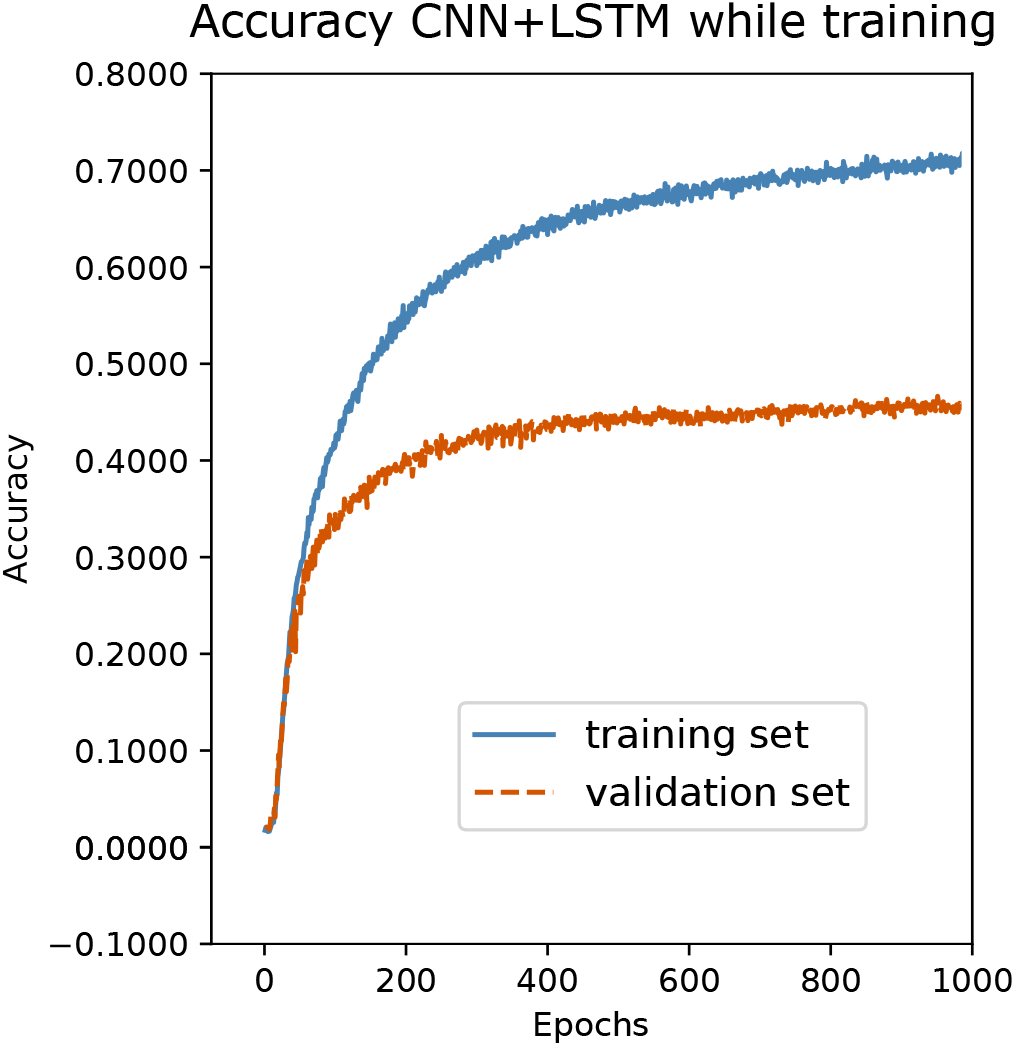
Training accuracy and validation accuracy over 1000 epochs on the influenza A dataset. With the CNN+LSTM architecture, the deep neural network shows no difficulties to learn.

**Figure S2:**
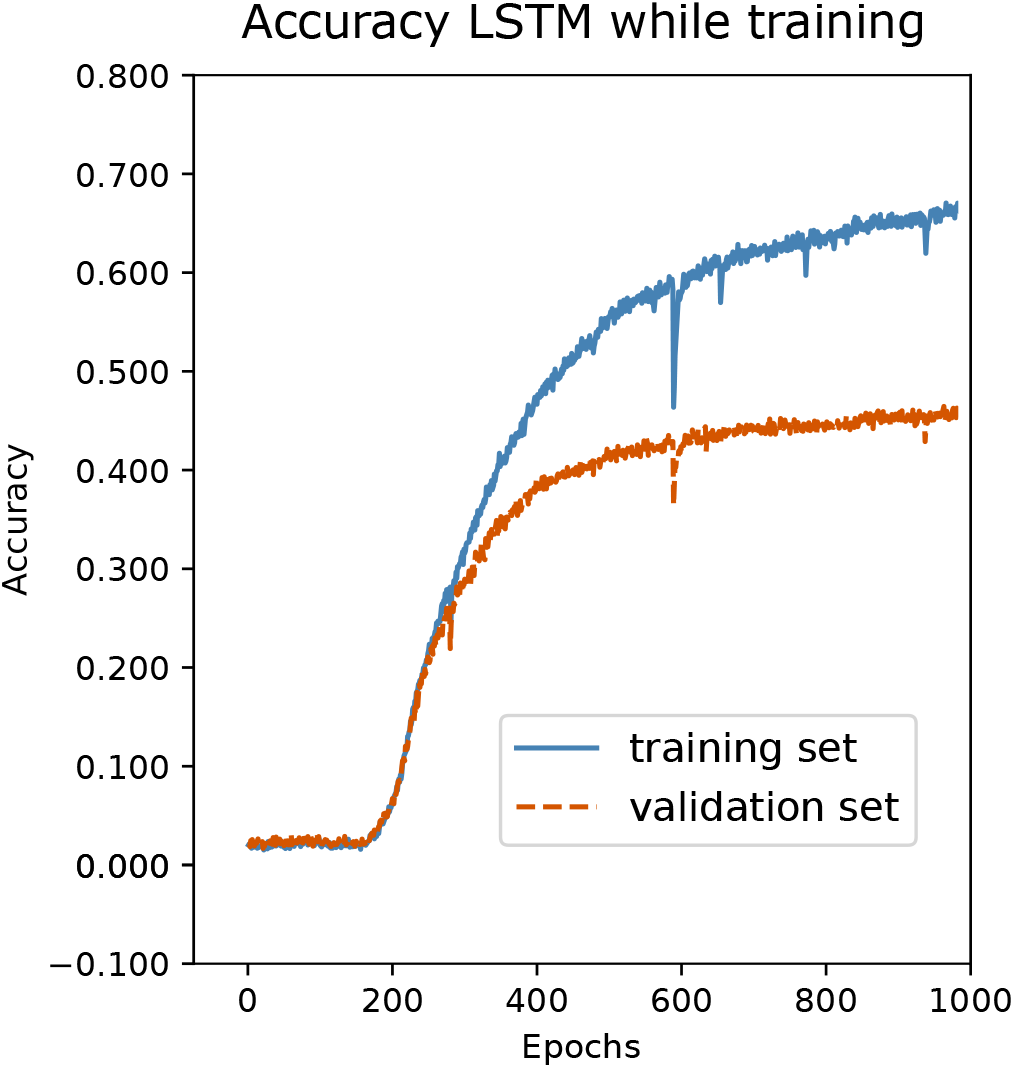
Training accuracy and validation accuracy over 1000 epochs on the influenza A dataset. With the LSTM architecture, the deep neural network remained in a state of random accuracy for ca. 200 epochs.

**Table S1.**
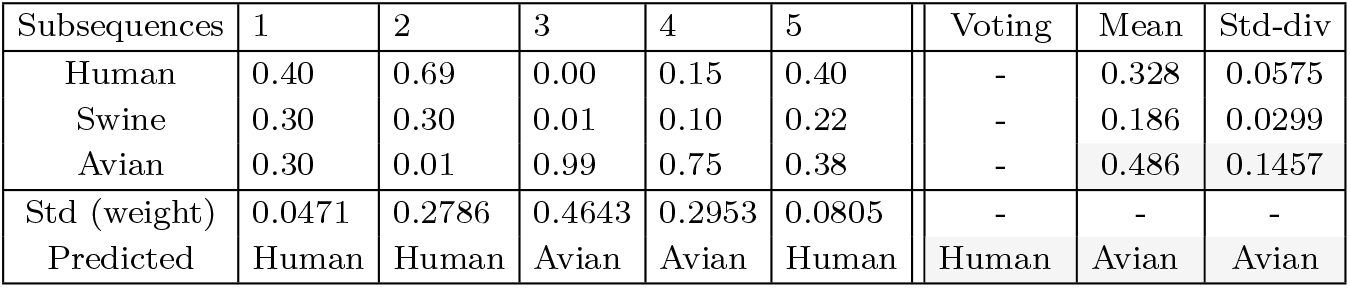
Comparison of host prediction results in regards to the different approaches that can be used to merge prediction scores of subsequences. In this example, a viral nucleotide sequence was split into five subsequences, and each of them was used to predict the corresponding host. Depending on the subsequence activation score merging approach, the final host prediction can vary.

**Table S2.**
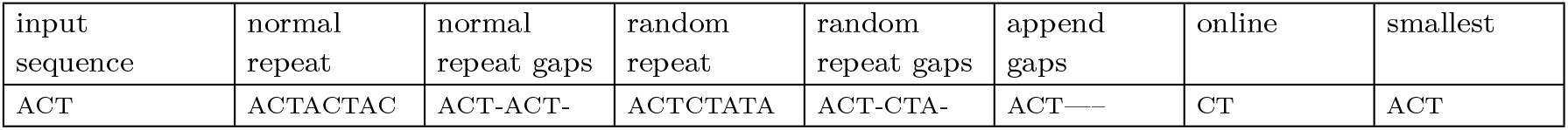
Comparison of the functional principle of the input expansion. Note that for real data, the raw data sequence would be hundreds of bases long. Furthermore, in this example, ACT is the shortest sequence of our dataset, but still longer than the input length expected by the neural network.

**Figure S3:**
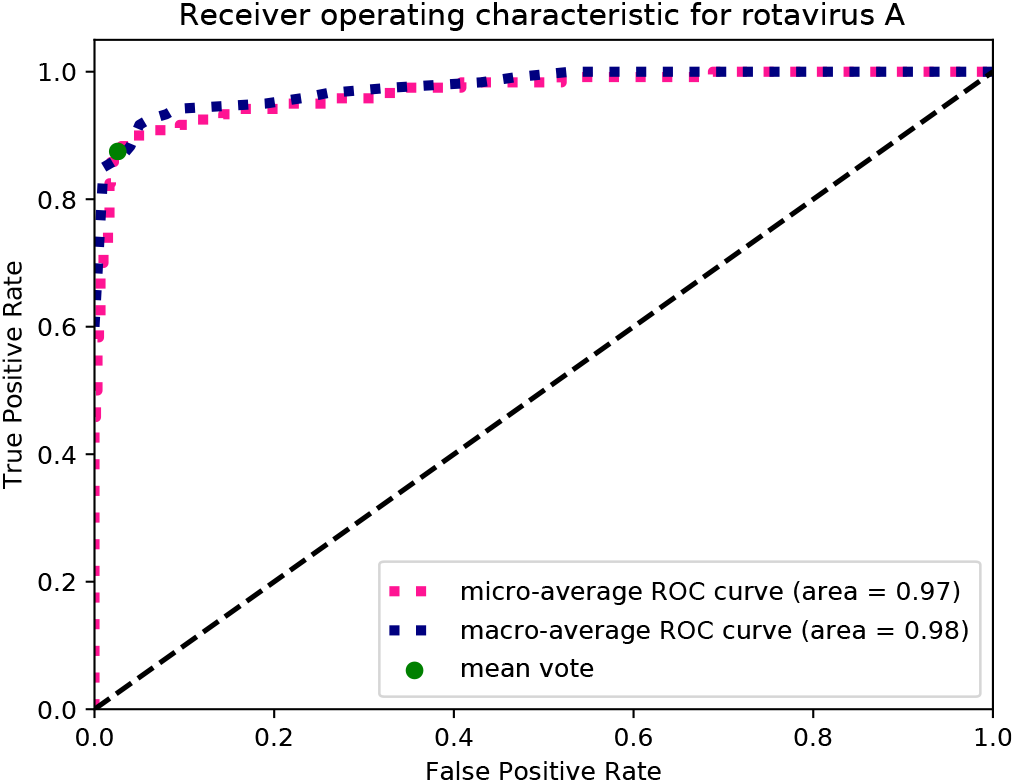
ROC curve for the rota dataset, calculated on the test set. The AUC of the micro-average ROC curve is 0.97 and for the macro-average 0.98. The trade-off between False Positive Rate and True Positive Rate, when using mean vote, is shown with a green dot.

**Figure S4:**
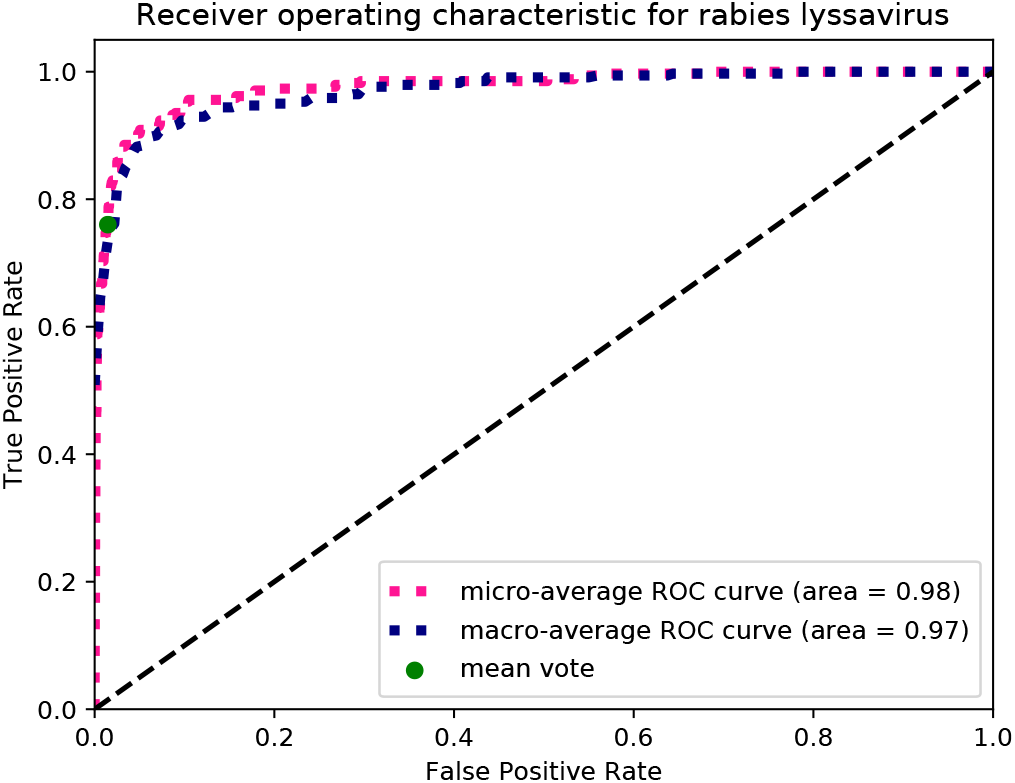
ROC curve for the rabies dataset, calculated on the test set.The AUC of the micro-average ROC curve is 0.98 and for the macro-average 0.97. The trade-off between False Positive Rate and True Positive Rate, when using mean vote, is shown with a green dot.

**Figure S5:**
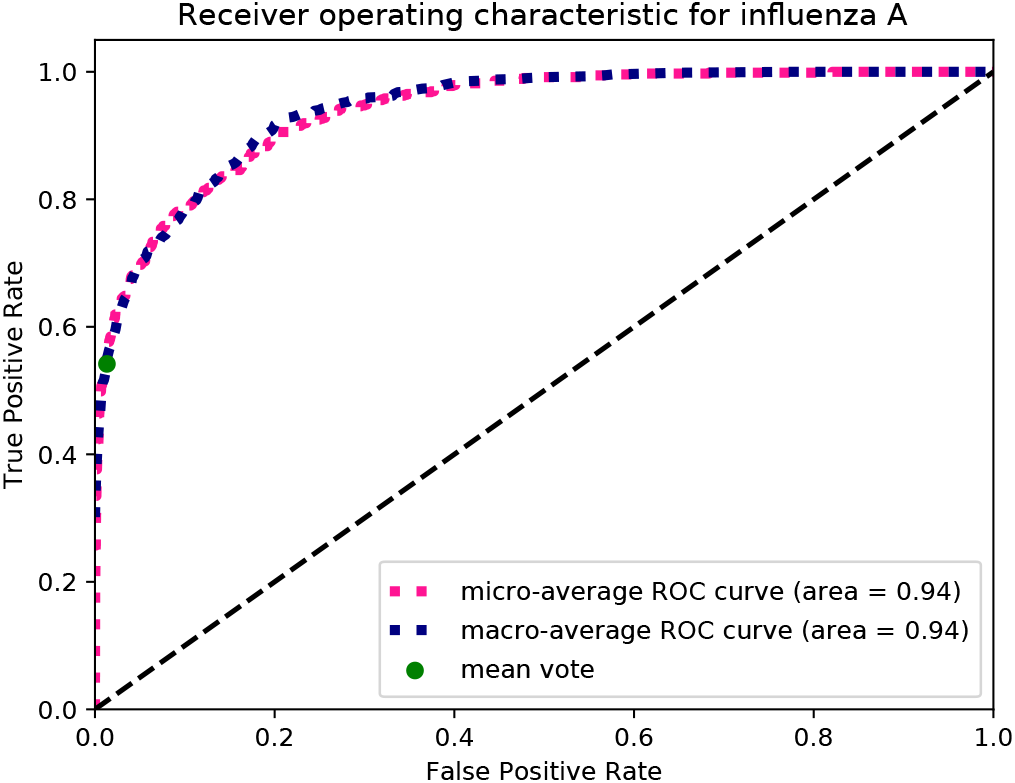
ROC curve for the influenza dataset, calculated on the test set. The AUC of the micro-average ROC curve is 0.94 and for the macro-average 0.94. The trade-off between False Positive Rate and True Positive Rate, when using mean vote, is shown with a green dot.

